# Phoneme-level processing in low-frequency cortical responses to speech explained by acoustic features

**DOI:** 10.1101/448134

**Authors:** Christoph Daube, Robin A. A. Ince, Joachim Gross

## Abstract

When we listen to speech, we have to make sense of a waveform of sound pressure. Hierarchical models of speech perception assume that before giving rise to its final semantic meaning, the signal is transformed into unknown intermediate neuronal representations. Classically, studies of such intermediate representations are guided by linguistically defined concepts such as phonemes. Here we argue that in order to arrive at an unbiased understanding of the mechanisms of speech comprehension, the focus should instead lie on representations obtained directly from the stimulus. We illustrate our view with a strongly data-driven analysis of a dataset of 24 young, healthy humans who listened to a narrative of one hour duration while their magnetoencephalogram (MEG) was recorded. We find that two recent results, a performance gain of an encoding model based on acoustic and annotated linguistic features over a model based on acoustic features alone as well as the decoding of subgroups of phonemes from phoneme-locked responses, can be explained with an encoding model entirely based on acoustic features. These acoustic features capitalise on acoustic edges and outperform Gabor-filtered spectrograms, features with the potential to describe the spectrotemporal characteristics of individual phonemes. We conclude that models of brain responses based on linguistic features can serve as excellent benchmarks. However, we put forward that linguistic concepts are better used when interpreting models, not when building them. In doing so, we find that the results of our analyses favour syllables over phonemes as candidate intermediate speech representations visible with fast non-invasive neuroimaging.

## Introduction

Speech perception is often conceptualised as a hierarchical process (Pisoni & Luce, 1987; DeWitt & Rauschecker, 2012). Concretely, the human brain is assumed to implement the extraction of semantic meaning from a highly dynamic sound pressure signal by means of a cascade of transformations into increasingly abstract representations. It is relatively clear that perceived speech sounds are first decomposed into a spectrally resolved representation at the cochlea. Facing greater challenges, further processing steps have been described for the various relay stations along the subcortical auditory pathway (Verhulst et al., 2018). Finally, there remains considerable uncertainty about the exact nature of cortical representations (Młynarski & McDermott, 2018).

One approach to gain insights into human speech processing is to employ encoding models. Here, the goal is to predict time-series of recorded neural data from the waveform of the stimulus presented during the recording. A popular framework organises this in two steps (Naselaris et al., 2011; Holdgraf et al., 2017): Nonlinear transformations of the stimulus material into various sets or spaces of features concretely capture hypotheses about cortical computations performed on the input signal, and a linear mapping of these feature spaces onto the neuronal responses then allows the evaluation of these hypotheses in terms of out-of-sample prediction performance. This way, data-rich naturalistic listening conditions of relatively long duration can be exploited, marking a considerable advantage in validity over isolated and artificial experimental paradigms (Theunissen & Elie, 2014; Hamilton & Huth, 2018). A range of recent results demonstrate the applicability of this approach across various neuroimaging modalities and research questions (Di Liberto et al., 2015; Huth et al., 2016; de Heer et al., 2017; Berezutskaya et al., 2017; Kell et al., 2018; Brodbeck et al., 2018).

A compelling finding obtained with this strategy is that predictions of cortical responses as measured with electroencephalography (EEG, Di Liberto et al., 2015) or functional magnetic resonance imaging (fMRI, de Heer et al., 2017) using acoustic feature spaces can be improved by additionally considering so-called articulatory feature spaces. The latter originate from the linguistic concept of representing a language with a set of smallest contrastive units, phonemes. Based on the observation that superior temporal regions are organised according to local selectivity to subgroups of phonemes, rather than individual phoneme identity (Mesgarani et al., 2014), the full set of phonemes is usually reduced by mapping each phoneme to its corresponding vocal gestures such as the voicing, tongue position or place and manner of articulation. Recently, it was shown that these manners of articulation are also readily decodable from EEG data time-locked to phoneme onsets in continuous speech stimuli (Khalighinejad et al., 2017). En- and decoding analyses based on articulatory feature spaces are thus interpreted as concordantly capturing a faculty which has been termed “pre-lexical abstraction” (Obleser & Eisner, 2008), i.e. a transformation of continuous physical properties of the waveform on categorical, invariant units of perception.

However, the transformation of speech stimuli into articulatory features comes with critical caveats. In addition to the original speech waveform, it requires a textual transcription thereof. This transcription must then be temporally aligned with the waveform and mapped to linguistically defined phonemes and corresponding articulatory features. A listener’s brain on the other hand has access to neither such a transcription nor linguistically defined mapping tables. Hence, the degree to which articulatory features specify a theory as to how the brain might compute such a representation is low. Models using articulatory feature spaces can thus be considered as *oracle models* (Kriegeskorte & Douglas, 2018). This leads to the question to which degree the gain in prediction in performance of articulatory features can be explained with features based on computationally more specified transformations of stimulus acoustics. Variance left unexplained by such acoustic features would suggest top-down signatures of a listener’s internal language model to be at play, affording the mapping of physical features of the input on a symbolic, categorical representation. Conversely, the extent to which directly processing the stimulus could account for the success of articulatory features would indicate how much of their predictive gain is actually attributable to bottom-up processing of the acoustic stimulus.

When choosing such acoustic feature spaces, one can proceed in different directions.

One possibility is that in order to explain the same variance as oracle models based on articulatory feature spaces, the detailed spectrotemporal characteristics that define subgroups of phonemes are needed. Correspondingly, one could progress from spectrograms to time series of activations of spectrotemporal patterns that have the potential to make information about relevant stimulus content such as distinctive phonemic characteristics more explicit while allowing to filter out irrelevant information about the original input. A physiologically inspired candidate feature space which interestingly has also been shown to improve the performance of automatic speech recognition software when used as input features (Schädler et al., 2012) are Gabor-filtered spectrograms. With this class of generic spectrotemporal kernels, one can describe several acoustic patterns that dissociate groups of phonemes. Examples are the spectral distance between formants as captured by filters of different spectral modulation, and formant transitions as captured by filters of joint spectrotemporal modulation. While this feature space has a long tradition in en- and decoding models of midbrain and auditory cortices in animal models and humans (Qiu et al., 2003; Pasley et al., 2012; Santoro et al., 2014, 2017; Berezutskaya et al., 2017; Holdgraf et al., 2017), it has so far never been applied to magneto- and encephalography (MEEG) data.

Another possibility is that the performance boost of the articulatory features is instead attributable to their correlation with a low-dimensional and less rich acoustic feature. It has repeatedly been observed that neuronal responses from bilateral superior temporal regions are especially sensitive to acoustic edges (Prendergast et al., 2010; Hertrich et al., 2012; Gross et al., 2013; Doelling et al., 2014; Hamilton et al., 2018; Oganian & Chang, 2018). Corresponding feature spaces extracting these onsets from envelope representations by means of half-wave rectifying the temporal gradient have previously been used by several labs (Hertrich et al., 2012; Hambrook & Tata, 2014; Fiedler et al., 2017). Additionally, features relying on the gradient capture the relationship of neighbouring time points, which is known to contain information about MEEG data across a range of different analyses (Ince et al., 2017). It is thus interesting to assess to which degree the gain in prediction performance obtained by articulatory features can be explained by such onset features.

**Figure 1:**
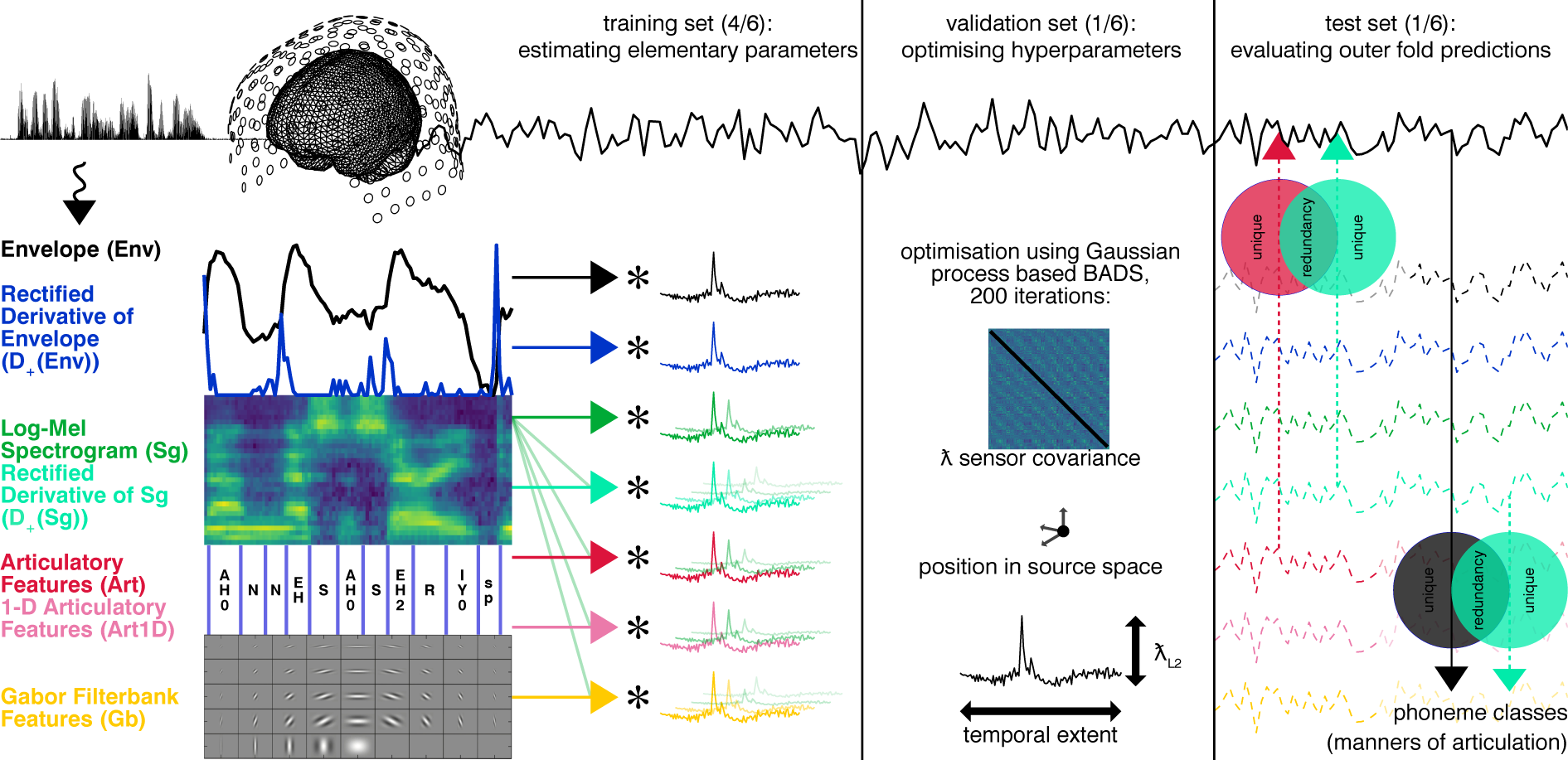
Concept of the study. Participants listened to a story of 1 hour duration while their MEG was recorded (top row). We nonlinearly transformed the speech waveform into various feature spaces (left column) and predicted neuronal responses with them using a linear finite impulse response (FIR) model (second column) in a nested cross-validation framework. We trained hyperparameters controlling the MEG source reconstruction as well as the FIR model (third column). We then evaluated the predicted responses of our encoding model by asking if a benchmark feature space using linguistic features was redundant with competing acoustic feature spaces using partial information decomposition (PID, right column, upward arrows). Additionally, we decoded four classes of phonemes from phoneme-locked observed MEG responses (right column, downward arrows). We then evaluated this decoding analysis by asking to what extent we could explain the same information from our encoding models, again using PID.

In the present study, we examine these two possibilities by comparing the predictive power of different acoustic feature spaces to that of an annotated articulatory feature space. We perform these investigations on an MEG story dataset of one hour duration per participant in a rigorously data-driven approach. A nested-cross validation framework (Varoquaux et al., 2017) is used to delegate the choice of model hyperparameters as well as MEG source level related preprocessing parameters such as dipole positions and sensor covariance matrix regularisation to a gaussian process based black-box optimisation algorithm (Acerbi & Ma, 2017). We thus allow encoding models based on different feature spaces the same chances to find optimal parameter combinations with a minimum of a-priori information while minimising the danger of overfitting. We then apply a recent implementation of partial information decomposition (PID, Ince, 2017) to assess to what degree the predictions of models based on acoustic features share information about observed recordings with predictions of the model based on articulatory features and to what degree these feature spaces contain information unique to them. Finally, the flexibility provided by this information theoretic framework allows us to quantify to what extent the information about manners of articulation decodable from phoneme evoked responses can be accounted for by predictions of our encoding models.

In this way, we aim to further elucidate to what degree we can non-invasively track signatures of an invariant processing stage related to acquired, internal knowledge of a language as opposed to a more simple reflection of bottom up encoding of physical properties of the stimulus.

## Results

### Correlations of responses to repeated chapters peak in bilateral auditory cortices

To identify regions in source space where MEG responses were repeatably activated by the stimulus (“story responsive” regions, Honey et al., 2012; de Heer et al., 2017), we correlated the responses to one chapter with the responses to its repetition. These correlations peaked in regions well in accordance with typical localisations of bilateral auditory cortices (AC, figure 2, top row).

**Figure 2:**
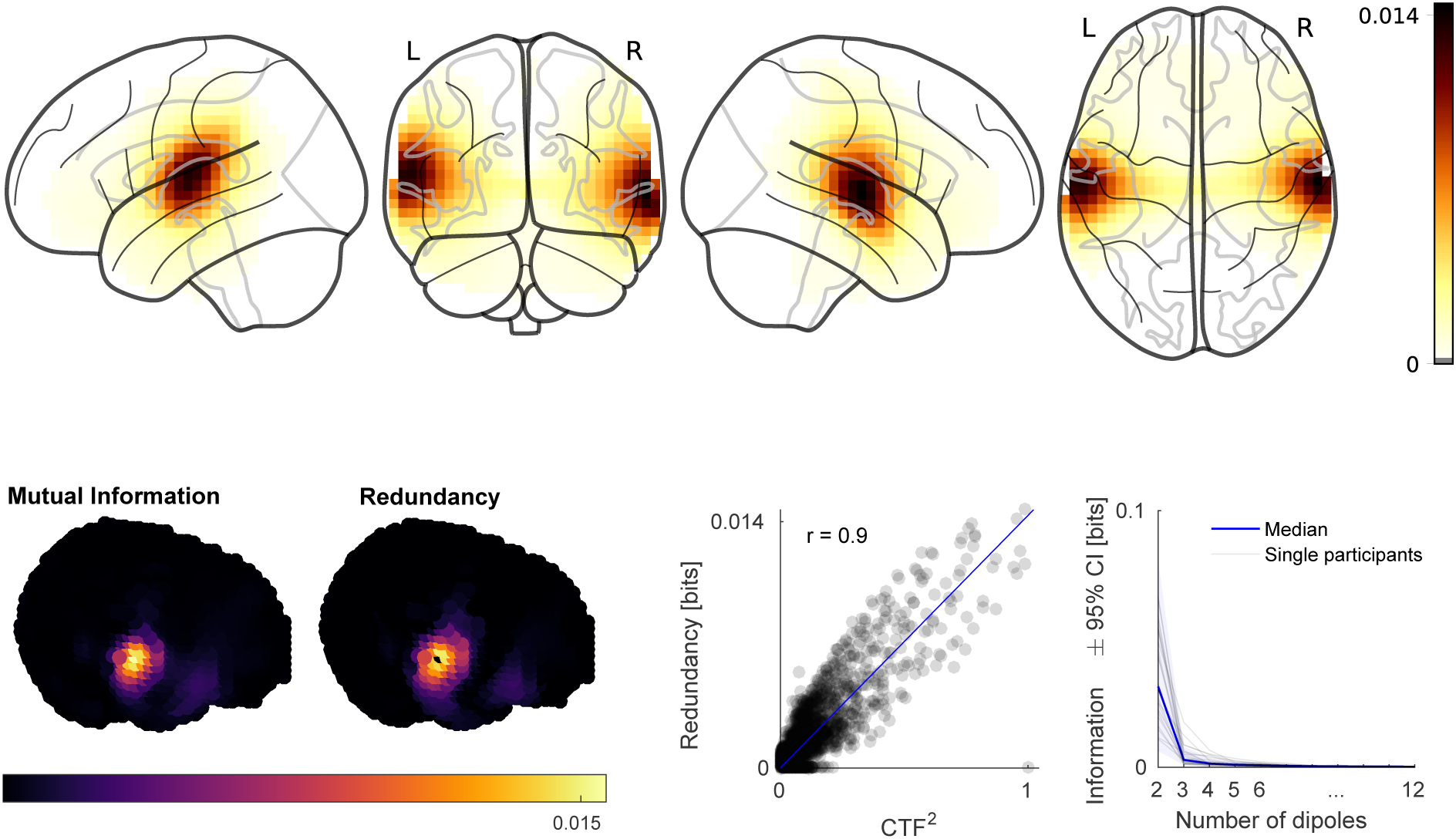
Identification and characterisation of story-responsive regions in source space. *Top:* Grand average of variance of brain activity of the last block explained by its repetition. *Bottom:* Left three plots show data of one exemplary participant. Left shows MI of activity in the first repetition about activity in the second repetition of the last block. Middle shows shared information (redundancy) of activity at bilateral story-responsivity peaks in the first repetition and activity in the first repetition at each other grid point about activity at each other grid point in the second repetition. Right scatterplot shows redundancy plotted against absolute values of the squared cross-talk function of the grid point of right peak story-responsivity, each dot is one grid point. Rightmost plot shows unique information added by sources additional to the bilateral story-responsivity peaks.

We then explored how much of the repeatable activity could be explained with how many dipoles in source space. To do so, we applied the framework of PID to the data of repeated blocks. PID (Ince, 2017; Williams & Beer, 2012) quantifies redundant (overlapping or common), unique and synergistic (super-additive) information of two source variables about a target variable (see for a more detailed description). Here, we were interested to what degree information in activity from the first repetition about activity in the second repetition at grid points different from the bilateral peaks of story-responsivity was shared with activity at the bilateral peaks. We found that maps of story-responsivity were well-matched by the maps of redundancy obtained by this analysis (figure 2, bottom row). Across participants, the average pearson correlation was .89 (averaging performed on fisher-z transformed correlations, then retransformed, range: .80 – .97). In most participants, this was to a large degree linearly attributable to the squared cross-talk functions (CTF) of the grid points of peak story-responsivity. Here, we found a mean correlation of .68 and .75 for the left and right hemispheres, respectively (ranges: -.09 – .94 and .26 – .91).

We then went on to iteratively identify grid points different from the bilateral peaks carrying unique information about the second repetition (see methods for a more detailed description). When we compared the unique information found across iterations to the information already available in bilateral peaks of story-responsivity, we found a characteristic elbow-pattern figure 2, bottom row, right plot). The median of the amount of unique information found in the grid point identified in the first iteration was below ten percent of the median of information already available in the bilateral peaks of story-responsivity. Based on these results, we focused the following analyses on one dipole per hemisphere, assuming that these would capture the essential part of the repeatable signal stemming from bilateral auditory cortices.

### Performance boost over spectrograms with annotated features, but also with acoustic features

The main goal of this study was to compare the predictive power of different feature spaces extracted from the speech stimulus about the identified relevant parts of MEG responses. For this, we trained (multidimensional) temporal response functions (mTRFs) using linear ridge regression. The elementary parameters of such models essentially result in FIR filters, one per dimension of a feature space. Additionally to these elementary parameters, we also optimised hyper-parameters of the models separately for each participant, hemisphere and feature space using a gaussian process based black-box optimisation algorithm (Acerbi & Ma, 2017). This way, we gave each feature space the same chances to optimally predict the MEG responses, avoiding a situation where using a fixed grid point across models might be biased towards a particular model due to the selection criteria (e.g. repeated block correlation). The hyper-parameters were the positions of one dipole per hemisphere and the amount of regularisation of the sensor covariance matrices of corresponding spatial filters as well as the amount of L2-regularisation of the ridge regression and the length or “temporal extent” of the FIR filters. The chosen positions converged to a spatially confined region in superior temporal gyrus (STG, figure S1, top) and did not exhibit systematic differences across feature spaces (figure S1, middle). The optimal sensor covariance matrix parameters were widely spread within the boundaries of the optimisation, tended to be larger in the left hemisphere and were relatively similar across feature spaces within the same participant (figure S1, bottom). The temporal extents of the learned filters (figure S2, top) as well as the amount of L2-regularisation (figure S2, bottom) exhibited characteristic values for the different feature spaces.

The performances of our models exhibited relatively large inter-participant variability and comparatively small variability across feature spaces (figure 3, top). To focus on systematic differences across feature spaces, we used Bayesian hierarchical linear modeling (Bürkner, 2017) and extracted the posterior distributions of betas of effects of the feature spaces. This allowed us to study the posteriors of differences between beta estimates for the benchmark feature space, a combination of spectrograms and articulatory features, and different competing feature spaces (figure 3, middle) as well as all other possible comparisons (figure 3, bottom left).

**Figure 3:**
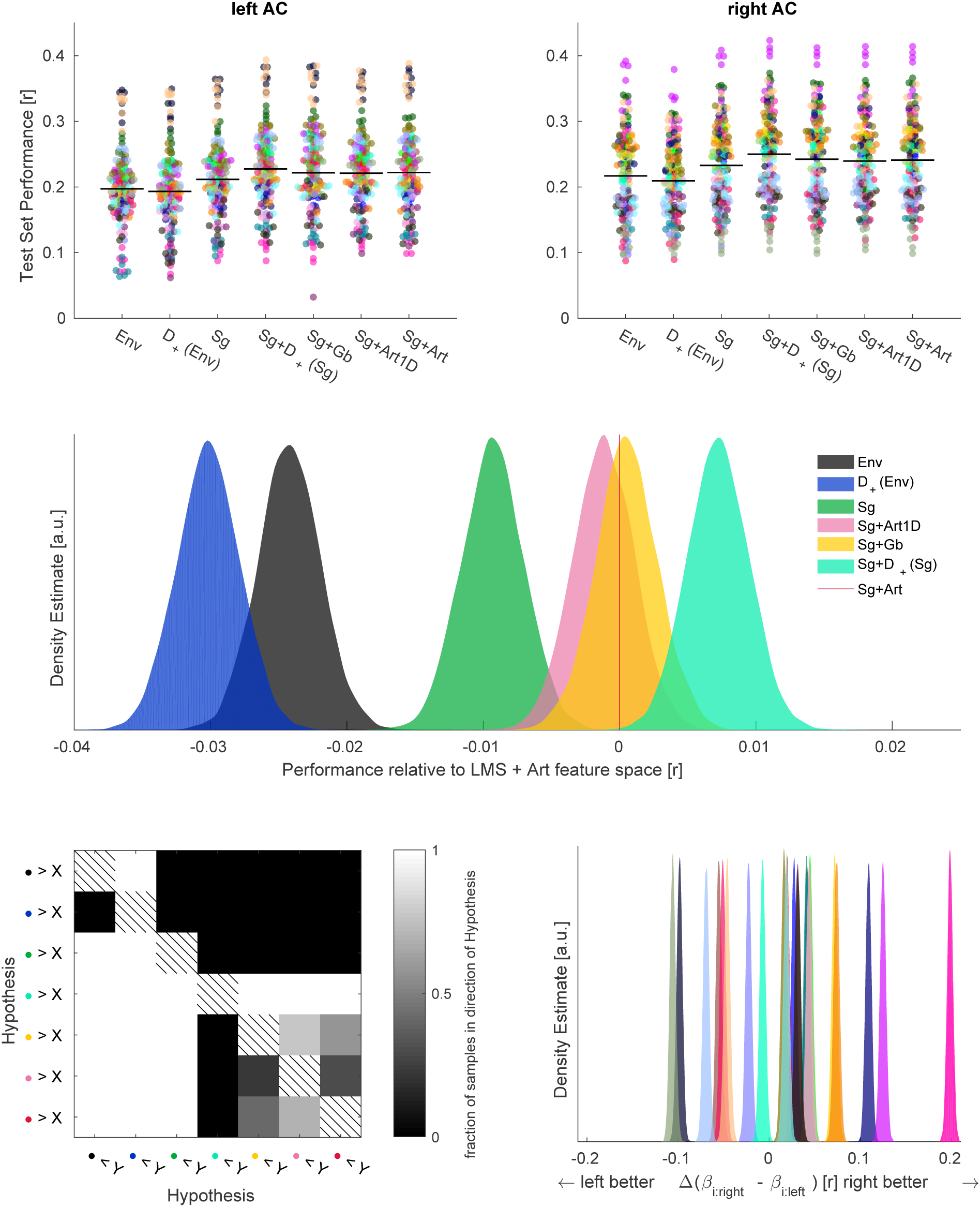
*Top:* Raw test set performances in left and right AC. Each colour is one participant, each dot is one outer fold. *Middle:* Samples from posterior distribution of differences of beta estimates of different feature spaces and Sg+Art feature space. *Bottom left:* Percent of samples in favour of hypotheses of differences of beta estimates between all feature spaces. Hypotheses are colour coded using the same colour mapping as in the middle plot, which corresponds to the bottom row and right column of the matrix shown here. *Bottom right:* Samples from posterior distribution of differences of beta estimates of individual participant’s right ACs minus left ACs. Colour mapping is the same as in the top plot.

We replicated previous results demonstrating an increase in prediction performance when combining linguistically motivated annotated articulatory features with spectrograms (*Sg+Art*, red vertical line) over spectrograms alone (*Sg*, fraction of samples of posterior distribution of differences in direction of hypothesis *f*_*h*_1__ = 1). When we discarded the pattern of articulatory features characteristic of each phoneme and thereby reduced the articulatory features to phoneme onsets (*Sg+Art1D*, pink), we obtained comparable performances (*f*_*h*_1__ = .99998). We also obtained comparable performances with a combination of spectrograms and Gabor-filtered spectrograms (*Sg+Gb*, yellow, *f*_*h*_1__ = .9998). However, this latter high-dimensional acoustic feature space was not outperformed by the combination of articulatory features and spectrograms (*f*_*h*_1__ = .54). Finally, we achieved the best prediction performances when we instead combined the spectrograms with their rectified temporal gradients (*Sg+D+(Sg)*, turquoise). This combination outperformed the combination of articulatory features and spectrograms (*f*_*h*_1__ = .99) as well as the combination of spectrograms and Gabor-filtered spectrograms (*f*_*h*_1__ = .98).

We also explored the lateralisation of the performances by evaluating within-participant differences of hemispheric betas that were independent from feature spaces (figure 3, bottom right). We found that the posterior distributions of differences of these hemispheric betas were narrow for single individuals, but exhibited a broad range of means within our sample: Some participants’ responses were easier to explain in left AC, some in right AC, while for others, there were no strong lateralisation effects.

These results raised the question whether the acoustic features explained the same or different aspects of the responses as the articulatory features.

### Performance boosts of articulatory features are highly redundant with those of a combination of spectrograms and their temporal gradient

PID Ince (2017); Williams & Beer (2012) aims to disentangle redundant, unique and synergistic contributions of two source variables about a target variable. Redundant information measures overlapping or common predictive power between the two sources about the target. Unique information is only available from one source, and synergistic information quantifies that part of the prediction of the target that can only be made when both sources are observed together on individual samples. We used the outer fold predicted MEG signal from two different models as the two source variables, and the corresponding recorded MEG signal as the target variable. This analysis therefore revealed how much the two feature spaces predicted the same parts of the MEG signal (redundancy), and how much each predicted distinct from the other (unique information). We fixed *Sg+Art* as one source, and considered the PID for the different acoustic feature spaces (figure4). We subjected the results to Bayesian hierarchical models similar to our analyses of the raw performances and focused on the betas of feature spaces.

**Figure 4:**
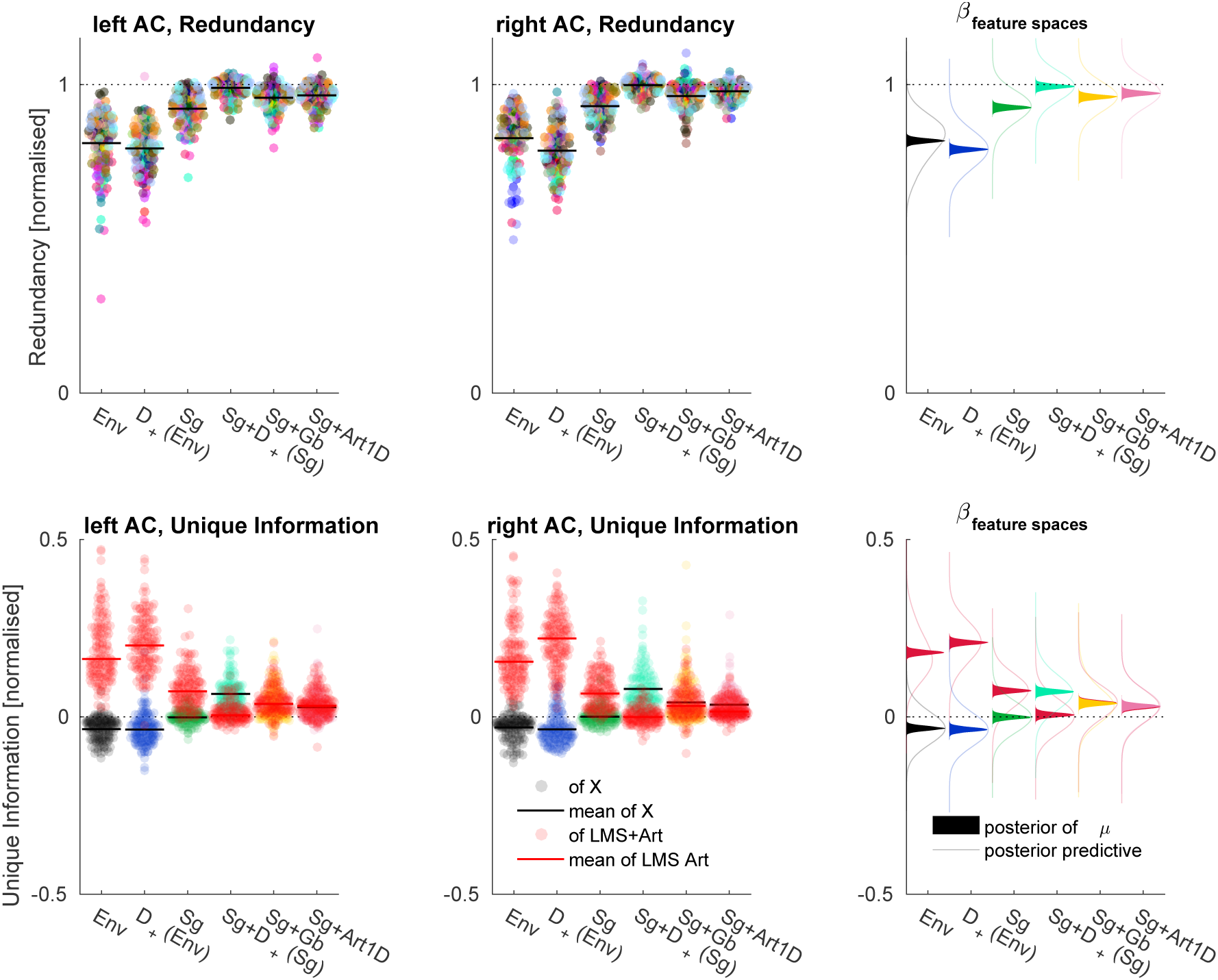
PID with test set predictions of articulatory vs other feature spaces as sources and observed MEG as target for left and right AC; values normalised by MI of prediction of articulatory features about observed MEG. *Top:* Redundancy; *Bottom:* Unique Information *Left and middle columns:* Raw results; each dot is one outer fold of one participant, pooled means overlaid; colour mapping corresponds to top row of figure 3. *Right column:* Solid lines show density estimates of posterior distributions of estimates of betas of feature spaces, transparent lines show density estimates of samples from posterior predictive distribution of the respective condition.

We observed a gain in redundancy of the multi-dimensional acoustic feature spaces such as *Sg* or *Sg+Gb* over the one-dimensional *Env* or *D*_+_*(Env)*, with all samples of posteriors of differences being in the direction in favour of the corresponding hypotheses (i.e. *f*_*h*_1__ = 1). This means, that the *Sg* and *Sg+Gb* feature spaces produce a prediction of the MEG signal which has more in common with the annotated *Sg+Art* model. The highest redundancy was achieved by *Sg+D* _+_*(Sg)*, reaching practically 100% of redundancy normalised by the information of the prediction from *Sg+Art* about the observed MEG (maximum a posteriori estimate (MAP) of corresponding beta: .9937, 95% credible interval (CI): .9795 – 1.0076, 95% CI of posterior predictive distribution: .8991 – 1.0862). It beat the second best acoustic model, *Sg+Gb* (*f*_*h*_1__ = 1) as well as the *Sg+Art1D* feature space (*f*_*h*_1__ = .9928).

Furthermore, we observed a reduction in information uniquely available from *Sg+Art* going from the one-dimensional to the multi-dimensional acoustic feature spaces (*f*_*h*_1__ = 1 for all corresponding hypotheses). For *Sg+D* _+_*(Sg)*, we even observed more unique information of the acoustic feature space than of the annotated feature space (*f*_*h*_1__ = 1), whose unique information was distributed around zero: The MAP estimate of the mean was .0056 (95% CI: −.0065 – .0181), and samples from the posterior predictive distribution had a 95% CI ranging from −.0888 – .0953. This suggests that all of the predictive power of the *Sg+Art* model is included in the *Sg+D_+_(Sg)* model. There is no unique information available in the *Sg+Art* prediction that a Bayesian optimal observer would not have been able to extract from the *Sg+D*_+_*(Sg)* model.

On a group level, these patterns were highly similar between left and right ACs.

### Model predictions fully capture phoneme-related characteristics of observed time-series

Another recently reported finding is that when epoching EEG data from a naturalistic story listening paradigm according to the phonemes constituting the stimulus, it is possible to decode four manners of articulation of these phonemes from the EEG data (Khalighinejad et al., 2017). We aimed to assess whether this was possible in our MEG data and to what degree our encoding models could account for this phenomenon.

To provide the decoding analysis with the best chances, we re-optimised the dipole position and sensor covariance matrix regularisation parameters of spatial filters. We did this as before using BADS (Acerbi & Ma, 2017), only this time with respect to the MI between MEG data epoched to phoneme onsets and the manner of articulation of each phoneme (4 discrete categories). In most cases, the positions found in this re-optimisation where very similar to those found before, however, in some cases, they differed slightly (figure S3, left). This MI was calculated separately for each time point in the extraced phoneme epochs. For the optimisation objective, we subsequently summed it across peri-phoneme onset time.

At the corresponding source locations, we found characteristic responses for the four manners of articulation (figure 5, top row, left: exemplary participant). We then retrained our encoding models based on all feature spaces with the source level parameters fixed to the values found when optimising for MI between MEG data epoched to phoneme onsets and the manners of articulation. In the outer fold predictions of these retrained models we observed phoneme-locked responses that were very similar to those obtained with observed MEG data (figure 5, top row, right).

**Figure 5:**
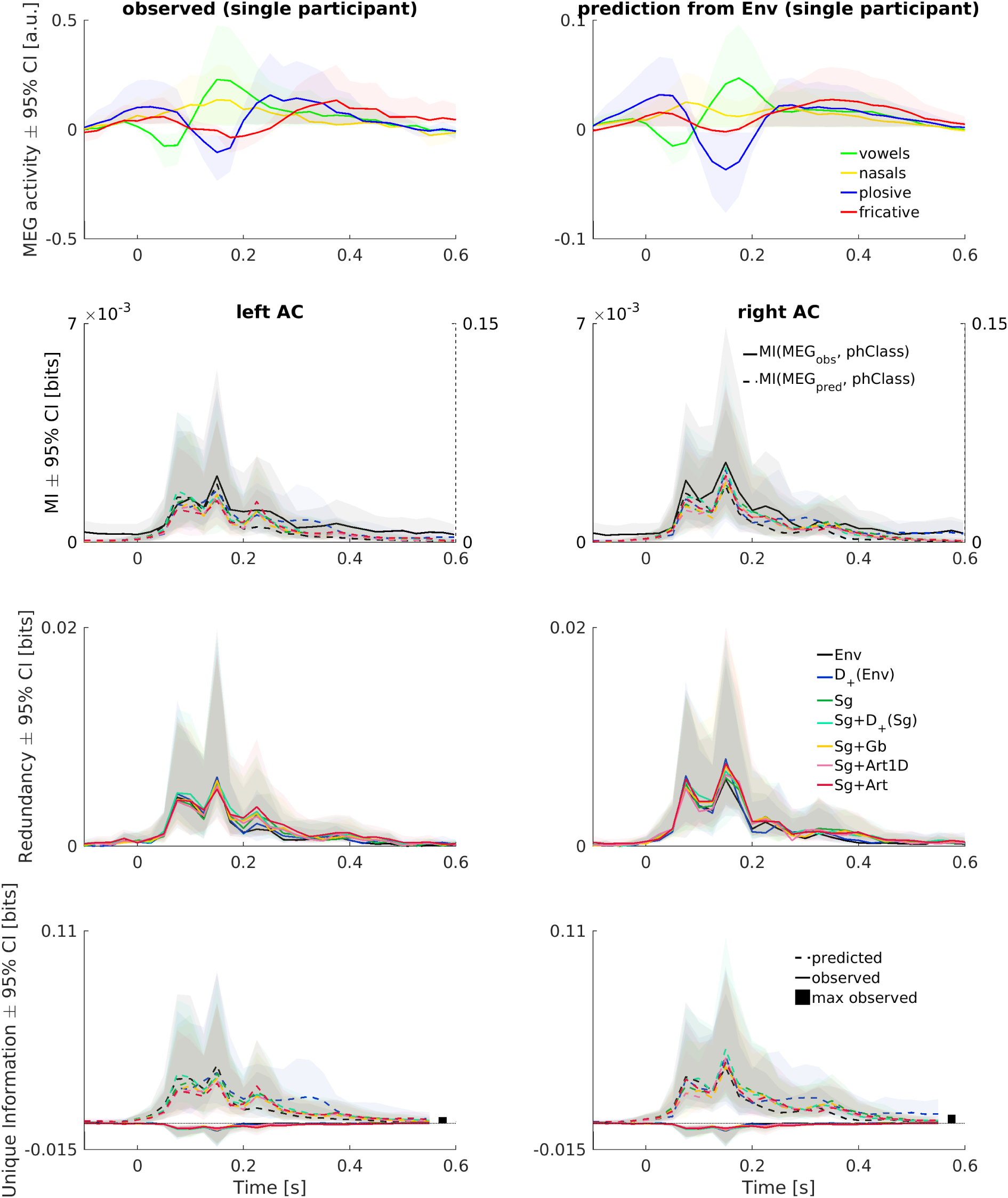
Phoneme related fields are captured by model predictions. *1st row:* Phoneme Related Fields of a single participant in observed (left) and predicted MEG (*Env* feature space, right). *2nd row:* MI of observed (solid lines, left y-axis) and predicted MEG (dashed and coloured, right y-axis) about four manner of articulation phoneme classes. *3rd row:* Redundancy from PID, amount of information that observed and predicted MEG share about manners of articulation. *4th row:* Unique Information of observed (solid) and predicted MEG about manners of articulation, maximum of information uniquely available from observed MEG across all participants, feature spaces and time points shown as black bars. Colour code in middle row applies to all rows. Rows 3 – 4 show medians across all participants *±* 95% (frequentist) confidence intervals, bootstrapped with 10000 samples.

Correspondingly, we observed a sustained pattern of MI following the phoneme onsets in bilateral ACs (figure 5, 2nd row, solid black lines, left y-axes). We found very similar patterns of MI between manners of articulation and predicted PRFs (dashed & coloured lines, right y-axes), with values roughly an order of magnitude higher. On a grand average level, this pattern of results did not substantially differ between either the two hemispheres or the different feature spaces.

To assess the amount of information shared by the observed and predicted time series about the manners of articulation as well as the amount of information unique to either of them, we performed PIDs with observed and predicted time series as sources and the manners of articulation as targets. This analysis could thus reveal if the observed MEG in fact contained information about manners of articulation different from that of e.g. the speech envelope convolved with a FIR filter.

The PIDs resulted in profiles of redundancy that very much resembled the marginal MI profiles for, on a group level, both hemispheres and all feature spaces alike (figure 5, 3rd row). Most importantly however, we also found that the information unique to the predicted MEG exhibited the same patterns (figure 5, bottom row, dashed lines). Crucially, the information unique to the observed MEG (solid lines) was negative, i.e. represented misinformation with respect to the other source, the predicted MEG. There was thus no relevant information about manners of articulation in the observed MEG that could not be retrieved from responses modeled with a convolution of any of our feature spaces with an FIR filter. Also here, this pattern of results was essentially the same for both hemispheres and all feature spaces.

Taken together, the results demonstrate that the models based on all of our feature spaces could fully account for the information about manners of articulation decodable from observed MEG responses.

## Discussion

We found that a model based on comparably simple acoustic features resulted in better prediction performance of MEG responses than a model based on annotated linguistic features. Furthermore, we found that predictions based on this best acoustic feature space were highly redundant with those of the linguistic feature space. This means that both models are providing the same prediction in the same time points. The amount of unique information was practically zero for the annotated feature space, but weakly positive for the best acoustic feature space. Finally, we showed that all of the feature spaces we considered here could fully account for the phonetic information decodable from phoneme-locked responses. In combination, these results show that a more parsimonious and accurate explanation for low-frequency MEG responses to continuous speech is provided by a bottom-up model of temporal stimulus dynamics rather than requiring higher-level linguistic constructs.

### Syllable level processing in superior temporal regions

In order to become consciously aware of the semantic meaning of the speech signal, the brain must eventually extract the necessary phonemic cues. The selectivity of high-gamma firing at spatially restricted regions in superior temporal gyrus to distinct phonetic features (Mesgarani et al., 2014) strongly suggests that such analyses are carried out in STG. Our results however discourage the view that signatures of this processing are observable in non-invasive, low-frequency time domain data. This begs the question what contribution the neuronal computations underlying these responses make to the extraction of meaning from speech input.

The acoustic edges which the MEG activity was found to respond to are classically related to a processing of speech on a syllable level (Greenberg et al., 2003; Hertrich et al., 2012; Gross et al., 2013; Doelling et al., 2014). Interestingly, in a recent ECoG study, it has been shown that similarly as observed here, also the high-gamma responses at regions in middle STG are best explained with features focusing on acoustic edges (Oganian & Chang, 2018). Further, the authors showed that in their stimulus material, the timing of these edges coincided with the transitions from syllable onsets to their central vowels, while their magnitude was found to correspond to the stress of syllables. As previously shown for low-frequency MEG data (Doelling et al., 2014), the high-gamma responses were independent of the phonemic content of the stimuli in that they also tracked acoustic edges in artificial non-speech stimuli. However, it is important to note that as we showed here, the reflection of the envelope already implies a reflection of distinctive manners of articulation (Khalighinejad et al., 2017), which are easily decodable from early stimulus representations (Nagamine et al., 2016). Taken together, this suggests that the low-frequency MEG responses described here are correlated with local neuronal firing which, independent of phonemic content beyond what is already explicit in the amplitude dynamics, coincides with the phonological structure of syllables.

At least from the perspective of low-frequency responses from STG to speech, the dominant role of the phoneme as mid-level speech features thus needs to be questioned. This is mirrored by a long tradition of linguistics, where various psychoacoustic phenomena and conceptual problems have been identified that are hard to explain with phonemes as the basic perceptual unit of speech (e.g. Massaro, 1974; Pisoni & Luce, 1987; Lotto & Holt, 2000; Holt & Lotto, 2010). A relatively recent flavour of reprimanding the dominance of phonemes comes from the success of applying differentiable programming (also referred to as “Representation -” or “Deep Learning”) to problems of natural speech processing. In this framework, mid-level features are the admittedly compelling byproduct of optimising weights connecting multilayered nodes in order to achieve a gradual compression of an initial input to a representation of maximal information relevant for a given task. The full potential of this technology is thus only exploited when handcrafted, theory-driven features are avoided and uncompromisingly data-driven end-to-end architectures can be implemented. Corresponding efforts yield state-of-the-art performance in speech recognition tasks (e.g. Deng et al., 2013) and can do so without access to phoneme dictionaries (Hannun et al., 2014). However, for many languages such as English, the character output of these ASR systems is necessarily highly correlated with a phonemic representation. When this forced mapping to phonemes is circumvented as in unsupervised dimensionality reduction architectures such as deep belief networks or autoencoders, a selectivity to phoneme-like features is not readily observable (Räsänen et al., 2016).

One way to instead demonstrate the computational significance of syllables is to reconsider the task the participants in the present study were given, which was to attentively listen to the story and understand its meaning in order to later answer questions about the content. Modern approaches in natural language processing allow to computationally describe the meaning of text by mapping its words into fixed dimensional embedding vectors that represent co-occurrence information of these words. When corresponding models (word2Vec or GloVe, Mikolov et al., 2013; Pennington et al., 2014) are trained on large text corpora, the resulting embedding vectors contain semantic information of the words of the employed dictionary. Interestingly, such a representation of words allows a prediction of activity of large parts of the human cortex as recorded by fMRI during story listening (Huth et al., 2016). Crucially, this principle has recently also been applied to speech instead of text corpora (Kamper et al., 2017; Chung & Glass, 2018), where an essential and non-trivial early step is to perform a segmentation on the speech material into word-like units. Despite currently suffering from relatively high word error rates, such unsupervised “zero-resource” speech models are an important step towards unbiased hypotheses about human speech recognition. Besides offering the fascinating perspective of making sense of unlabeled speech data, they are thus ideally suited for a fruitful mutual exchange of research in the domains of artificial intelligence and cognitive neuroscience. For example, it has been found that suitable segmentation algorithms benefit from incorporating the classic neurophysiological model (Ghitza, 2013) of an oscillatory segmentation process (Räsänen et al., 2015).

Another perspective on the responsiveness of the auditory cortex to acoustic edges is provided by automatic speech recognition systems which aim to transform spoken language into textual representations. Here, it has been found that features very similar to our winning feature space could increase the robustness to noise and reverberation (Kumar et al., 2011). From that perspective, it is conceivable that the observed relationship between stimulus energy and neuronal dynamics is indeed reflecting an optimisation for adverse listening conditions, arguably one of the hardest and ubiquitous challenges for the auditory system. Focussing on the rapid power changes in the speech signal then is an efficient way to suppress slowly-changing, stationary types of noise. However, deeper insights into suitable neuronal coping strategies are better gained with studies sampling the stimulus space correspondingly (Giordano et al., 2016; Fiedler et al., 2018), not with the present dataset restricted to clear speech.

### Hyperparameter optimisation as a principled approach to parameter settings

Given the relatively small amount of variance attributable to the different feature spaces and the comparably large amount of variance across participants, it is important to note that for applications based on this family of forward models (Mirkovic et al., 2015; Fiedler et al., 2017), it might be of less relevance to focus on the feature spaces considered here. The lion’s share of predictive power was achieved with the by now classical speech envelope, a finding in line with the effect sizes reported in a recent application-oriented and EEG-based comparison of acoustical feature spaces (Biesmans et al., 2017). The model performances we report here are however considerably higher than those reported in other studies employing linear encoding models on non-invasive MEEG data (Di Liberto et al., 2015; Wong et al., 2018). While we do not know how the individual improvements in our modeling pipeline contributed to this increase in performance, we attribute it mainly to our model’s spatial and temporal specificity to the neuronal response characteristics of individual participants and feature spaces. This was made possible by including a data-driven and efficient hyperparameter optimisation algorithm (Acerbi & Ma, 2017) into our pipeline.

Given the typically low Signal-to-Noise ratios (SNRs) in neuroimaging, an optimisation of analysis parameters is necessary. Traditionally, such an optimisation entails a dramatic increase of the ‘researcher degrees of freedom’. Usually, such a strategy is thus discouraged as it increases the danger of overfitting and easily compromises the reproducibility of results (Nichols et al., 2017). However, the use of a robust nested cross-validation framework (Varoquaux et al., 2017) ensured that our final model evaluations were performed on test data unseen while independently optimising both elementary and hyperparameters on training- and validation sets respectively. We thus put forward that such a strategy allows a significant reduction in bias in the exploitation of large data sets such as ours.

The benefits of this approach are manifold. Most prominently, it is possible to evolve from parameter settings that are traditional in the field to potentially surprising optimal choices.

In the present study, the optimal choices found for the temporal extent of the encoding models – the length of the learned filters – for different feature spaces are considerably longer than previously used (Ding & Simon, 2012; Di Liberto et al., 2015; Brodbeck et al., 2018). Also, optimal parameters revealed characteristic ranges for different feature (sub-)spaces, showcasing a flexibility that can adapt to detailed parameter patterns across and within participants. In datasets where multiple regions of interest with similar signal-to-noise ratios can be identified, it is in theory also possible to compare the temporal extents of these regions (Lerner et al., 2011). Such an analysis would be conceptually similar to the characterisation of spatial “feature pooling” as recently implemented by St-Yves & Naselaris (2017).

Another hyperparameter whose optimal choices substantially differed from those traditionally applied in the field was the regularisation of the sensor covariance matrices. Data-driven approaches to this problem have been suggested previously (Woolrich et al., 2011; Engemann & Gramfort, 2015). The rationale here differs in its optimisation objective. We were searching for a parameter value that is low enough to prevent a mixing of source estimates with unrelated sources while being high enough to provide a robust, noise suppressive spatial filter in order to maximise the performance of the encoding model. The optimal values we found then were often considerably higher than the usually chosen 5%. Moreover, while exhibiting a large degree of stability across the different feature spaces within a hemisphere of a given participant, a lot of variability was found not only across participants but also across hemispheres, likely reflecting drastically different signal-to-noise ratios in the respective situations.

An additional benefit of the hyperparameter optimisation as performed here is that it allowed us to treat the hyperparameters as continuous, in contrast to what is necessary for grid search approaches. While becoming computationally infeasible with higher numbers of hyperparameters, it also requires a discretisation of the parameters. This is traditionally done for the regularisation in the ridge regression models (Crosse et al., 2016). Another example is the use of grids in MEG source space. With the computing power available at present, it has become a standard approach to sample the source space with a cubic grid and perform “whole-brain” mass-univariate analyses. While this is an indispensable tool, e.g. for initial data exploration as in our case, a spatial position of maximum effect can only be found with a high resolution of the grid, which however often also drastically increases the number of redundant grid points (Faharibozorg et al., 2018) and thus renders computationally demanding analyses cumbersome to run. In our current approach, we treated the source position coordinates as a continuous parameter and navigated to optimal positions within the usually smooth and thus easily optimiseable space in the volumes. A similar approach has recently also been developed (Lorenz et al., 2016) and applied (Lorenz et al., 2018) for closed-loop applications in fMRI.

For our present study, it allowed us to rule out that our findings were biased towards low-level processing. This would possibly have been the case with a more pragmatic focus on grid points of peak repeatability, as it is possible that participants paid less attention to the second presentation of the last block. Instead using these peaks as initial values of our optimisation alleviated the identification of superior temporal regions, but critically allowed us to fine-tune the exact position. While it did not find consistently different source positions for different feature spaces as it might have been expected given results from other imaging modalities (DeWitt & Rauschecker, 2012; Hamilton et al., 2018), the region across which these source positions were clustered was relatively confined, evidencing the convergence of the black box optimisation algorithm.

Finally, the spatially confined source position estimates also corroborate our assumptions about the origin of our signal: Similarly to previous efforts (Ding & Simon, 2012), we restricted ourselves to a single dipole per hemisphere, since further dipoles were highly redundant and contributed only a small amount of unique information. This reduction in dimensionality allowed us to run our extensive optimisation procedure individually for each point in source space, which is an improvement over choosing parameters to be optimal for an average of activity from many regions. Since in our case this would most likely include many regions with unrelated activity, parameters as for example those controlling the temporal extent would converge on values underfitting the actual signal in order to best fit a mixture of signal and noise.

The approach can thus be seen as a powerful option to identify functional regions of interest and hence adds to the toolkit of data-driven, bottom-up analyses of MEG data. As the overall framework thus not only selects suitable stimulus features but also operates on the neuronal responses, it can in principle be seen as similar to recent suggestions based on Canonical Component Analysis (CCA, de Cheveigné et al., 2018). Here however, we maximise the anatomical interpretability of the resulting spatial filters by design, circumventing a problem inherent to CCA.

Taken together, setting model and preprocessing related hyperparameters by means of a black-box optimisation algorithm constitutes a relatively easily implementable and flexible way of realising an adaptive model. With greater computational efforts, i.e. more samples from the hyperparameter space, it is in principle also possible to extend the rationale to even more preprocessing parameters (Hahn et al., 2018). The present approach thus marks an important step towards more principled and data-driven pipelines in neuroimaging (Bzdok & Yeo, 2017).

### Conclusion

In a rigorously data-driven approach, we have studied models that explain cortical neuronal responses as captured by source-localised MEG in a story-listening paradigm. Our results underscore that annotated linguistic feature spaces are useful tools to explore neuronal responses to speech and serve as excellent benchmarks. We find their performance for explaining neuronal responses of high temporal resolution to be exceeded and explained by a comparably simple acoustic feature space considering spectrotemporal dynamics and thus conclude that a more parsimonious explanation of the gain in predictive performance over spectrograms alone rests on onset sensitivity of the auditory system.

## Methods

### MEG recording, preprocessing and spatial filtering

#### MEG recording

24 healthy young participants (native speakers of English, 12 female, mean age 24.0 years, age range 18 – 35 years) agreed to take part in our experiment. They provided informed written consent and received a monetary compensation of £9 per hour. The study was approved by the College of Science and Engineering Ethics Committee at the University of Glasgow (application number: 300170024). Participants listened to a narrative of 55 minutes duration (“The Curious Case of Benjamin Button”, public domain recording by Don W. Jenkins, librivox.org) while their brain activity was recorded with a 248 channel magnetometer MEG system (MAGNES 3600 WH, 4D Neuroimaging) at a sampling rate of 1017.25 Hz (first 10 participants) and 2034.51 Hz (last 14 participants). Prior to recording, we digitised each participant’s headshape and attached five head position measurement coils to the left and right pre-auricular points as well as to three positions spread across the forehead. The session was split into 6 blocks of equal duration and additionally included the repetition of the last block. The last ten seconds of each block were repeated as a lead-in to the following block to allow listeners to pick up the story. Prior to and after each block, we measured the positions of the coils. If the movement of any of them exceeded 5 mm, we repeated the block. After the recording, participants had to answer 18 multiple choice questions with 3 options each, where the number of correct options could vary between 1 and 3 per question. The questions referred to the entire story, covering three details per recording block. The average performance was .95 with a standard deviation of .05 and a range from .78 to 1.00.

#### MEG preprocessing

Most of our analyses were carried out within the Matlab computing environment (v2016a, MathWorks, Natick, MA, USA) using several open-source toolboxes and custom code. Deviations from this are highlighted. Preprocessing was done using the fieldTrip toolbox (Oostenveld et al., 2011). Initially, we epoched the data according to the onsets of the full blocks including the ten seconds of lead-in. For noise cancellation, we subtracted the projection of the raw data on an orthogonal basis of the reference channels from the raw data. We manually removed and subsequently replaced artefactual channels with spherical spline interpolations of surrounding channels (mean number of artefactual channels per block: 3.07, standard deviation: 3.64; pooled across participants), replaced squid jumps with DC patches, filtered the signal with a fourth-order zero-phase butterworth high-pass filter with a cutoff-frequency of .5 Hz and downsampled the data to 125 Hz. We then excluded the lead-in parts from the blocks and performed Independent Component Analysis (ICA) to identify and remove components reflecting eye and heart activity (mean number of components per block: 6.70, standard deviation: 5.01; pooled across participants) and further downsampled the data to 40 Hz.

#### MEG source space

We employed three different source modeling approaches for our analysis. Firstly, we aimed to identify regions in source space whose activity was in a repeatable relationship with our auditory stimulation (”story-responsive” regions, de Heer et al., 2017; Honey et al., 2012). Secondly, we wished to visualise these results on a group-level. Lastly, for our main intention of modeling the story-responsive regions, we designed a framework that would allow us to optimise parameters of our spatial filters as part of a cross-validation, similar to a recent proposal by Engemann & Gramfort (2015).

##### Volume conductor models

For all three approaches, we obtained common volume conductor models. We first aligned individual T1-weighted anatomical MRI scans with the digitised headshapes using the iterative closest point algorithm. Then, we segmented the MRI scans and generated corrected-sphere volume conductor models (Nolte, 2003). We generated grids of points in individual volumes of 5 mm resolution. For group-level visualisation purposes, we also generated a grid with 5 mm point spacing in MNI space, and transformed this to individual spaces by applying the inverse of the transform of individual anatomies to MNI space.

##### Initial data exploration: identification and characterisation of story-responsive regions

To identify story-responsive regions in MEG source space, we projected the time-domain sensor level data through rank-reduced linearly constrained minimum variance beamformer spatial filters (Van Veen et al., 1997) with the regularisation of the sensor covariance matrices λ*_s_* set to 5%, using the dipole orientation of maximal power. We correlated the responses to the last block with those to its repeated presentation within each participant to obtain maps of test-retest-*R*^2^ (“story-responsive” grid points, Honey et al., 2012; de Heer et al., 2017). We repeated this using the grids in MNI space we had warped into individual anatomies for a group-level visualisation using the *plot_glass_brain* function of the Python module Nilearn (Abraham et al., 2014).

We then explored how many dipoles would explain how much of the repeatable activity in story-responsive regions. It is known that due to the non-uniqueness of the inverse problem, the spatial resolution of MEG source reconstructions is inherently limited. Neighbouring grid points are thus often highly correlated, rendering analyses on a full grid highly redundant (Faharibozorg et al., 2018). To avoid such an unnecessary computational burden for our modeling, we used an information theoretic approach to characterise redundant and unique regions in source space.

First, we computed Mutual Information (MI, Ince et al., 2017) at each grid point in individual source spaces between activity in the first and the second repetition, essentially repeating the initial identification of story-responsive regions. Next, we applied the framework of partial information decomposition (PID, Ince, 2017) to the data of repeated blocks in an iterative approach. PID (Ince, 2017) aims to disentangle redundant, unique and synergistic contributions of two source variables about a target variable. As the first source variable, we here used the two-dimensional activity at bilateral grid points of individual peak story-responsivity during the first repetition. We then scanned the whole grid in parallel for both repetitions, using the activity recorded during the first repetition as the second source variable and the activity recorded during the second repetition as the target variable of PIDs. We were then interested in the resulting maps of redundancy and unique information. The former would allow us to infer to what degree other grid points with high story-responsivity shared their information about the repetition with the grid points of peak story-responsivity. The latter on the other hand would show us where information unexplainable by these two peaks could be found. After this first iteration, we added the grid point of peak unique information to the then three-dimensional first source variable in the PIDs and repeated the computation across the whole grid. We reran this approach for a total of ten iterations. Finally, we computed MI between the two-dimensional activity at bilateral peaks of story-responsivity in the first and the second repetition and compared this to the unique information found in each iteration of our iterative approach.

##### Optimisation of source space coordinates and sensor covariance regularisation

In order not to unnecessarily spend computational resources, we wanted to limit our main endeavour of modeling MEG responses to parts of the signal which actually were in a systematic relationship with the stimulus. A straight-forward solution for a selection of these parts would have been to directly use the grid points identified as story-responsive using the test-retest correlation. However, since it is likely that participants paid a lesser degree of attention to the more predictable repeated presentation of the last chapter, we could not rule out that the test-retest-*R*^2^ maps would be biased towards low-level auditory processing. Furthermore, these maps could be influenced by differences in the position of the participant’s head in the scanner as well as the amount of eye blinks and head movements. The peak test-retest points are thus not guaranteed to be the optimal locations for any given feature space model fit and tested over the whole experiment. Moreover, it was possible that different feature spaces would optimally predict distinct regions. Finally, we did not know a-priori what level of regularisation of the sensor covariance matrices would be ideal to capture the responses of interest in each individual dataset.

To account for all of these considerations in a data-driven manner, we treated the coordinates of regions of interest as well as the regularisation of sensor covariance matrices as hyperparameters of our model, which we optimised by means of a black-box optimisation algorithm. We kept the other specifications of the spatial filter design as described above. As initial coordinates, we used the maxima of test-retest-*R*^2^ maps within each hemisphere. The boundaries of the coordinate hyperparameters were defined by the boundaries of the respective hemisphere of the individual brain volume which we shrunk by a factor of .99 for this purpose to avoid instabilities of the forward models close to their boundaries (table 1). In each iteration of the black-box optimisation, we then applied a given amount of regularisation to the precomputed sensor covariance matrix and computed the leadfield for a given vector 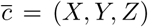 of coordinates in source space using the precomputed volume conductor model for each block. Since the orientation of the resulting dipoles was then arbitrary, i.e. possibly flipped across blocks, we estimated the mean axis of dipoles across blocks and changed the sign of the orientations of dipoles whose dot products with the orientation of the dipole closest to the mean axis were negative. We then recomputed the leadfields for these aligned dipole orientations. Finally, we projected the sensor level data through these spatial filters and z-scored them within each block to account for differences in mean amplitude across blocks.

**Table 1:**
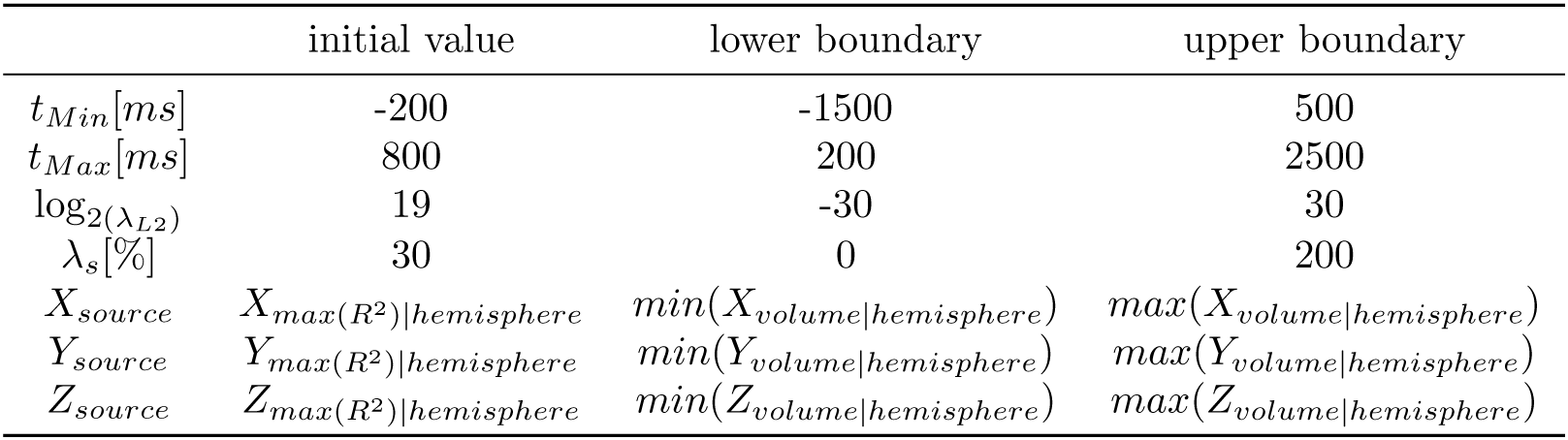
Initial values and boundaries for hyperparameters in BADS optimisation.

|  | initial value | lower boundary | upper boundary |
| --- | --- | --- | --- |
| $t_{Min}[ms]$ | -200 | -1500 | 500 |
| $t_{Max}[ms]$ | 800 | 200 | 2500 |
| $\log_2(\lambda_{L2})$ | 19 | -30 | 30 |
| $\lambda_s[\%]$ | 30 | 0 | 200 |
| $X_{source}$ | $X_{max(R^2)\vert hemisphere}$ | $min(X_{volume\vert hemisphere})$ | $max(X_{volume\vert hemisphere})$ |
| $Y_{source}$ | $Y_{max(R^2)\vert hemisphere}$ | $min(Y_{volume\vert hemisphere})$ | $max(Y_{volume\vert hemisphere})$ |
| $Z_{source}$ | $Z_{max(R^2)\vert hemisphere}$ | $min(Z_{volume\vert hemisphere})$ | $max(Z_{volume\vert hemisphere})$ |

##### Evaluation of distinctiveness of optimised coordinates for different feature spaces

The main results of this optimisation were triplets of coordinates in source space for each outer fold of each feature space in each hemisphere of each participant. To evaluate the degree to which these source coordinates were distinct for different feature spaces, we computed the silhouette index. As a measure of the consistency of a clustering, it relates the similarity of data within a given class to the similarity of data outside of that given class and is bound between −1 and +1. For the optimised source positions of each outer fold *o* of the set of outer folds *O* and each feature space *f* of the set of feature spaces *F*, we computed the silhouette index *s* (*o_f_*) using the following formula:

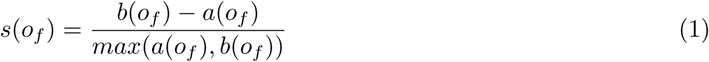

Here, *a*(*o_f_*) denotes the average euclidean distance between the source position chosen in the outer fold *o* and the source positions chosen in *O \ o* for that feature space *f*, while *b*(*o_f_*) refers to the minimum of average distances between the source position chosen in the outer fold *o* for feature space *f* and source positions chosen for all outer folds in *O* for all feature spaces in *F \ f*.

### Stimulus transformations

The speech stimulus was then transformed into various feature spaces. We used the GBFB toolbox (Schädler et al., 2012) to obtain 31-channel Log-Mel-Spectrograms (*Sg*, ranging from 124.1 Hz – 7284.1 Hz) and summed these across the spectral dimension to also obtain the amplitude envelope (*Env*). Additionally, we filtered the spectrograms with a set of 4552D Gabor filters of varying centre frequencies corresponding to those of the *Sg* as well as spectral modulation frequencies (0, 2.9, 6 12.2 and 25 Hz) and temporal modulation frequencies Ω (0, 6.2, 9.9, 15.7 and 25 Hz, “*Gb*”). Notably, this implementation of the toolbox only considers a subset of all possible combinations of centre frequency as well as spectral and temporal modulation frequencies to avoid overly redundant features. As a last acoustic feature space, we computed half-wave rectified first derivatives of the envelope and spectrogram feature spaces (“*D*_+_*(Env)*” and “*D*_+_*(Sg)*” Hertrich et al., 2012).

To construct annotated feature spaces, we used the Penn Phonetics Lab Forced Aligner (Yuan & Liberman, 2008) to align the text material to the stimulus waveforms, providing us with onset times of phonemes comprising the text. These were manually corrected using Praat (Boersma, 2001) and subsequently transformed into a 23-dimensional binary articulatory feature space (“*Art*”, de Heer et al., 2017). Finally, we discarded the information about phoneme identity to obtain a one-dimensional binary feature space of phoneme onsets (“*Art1D*”).

Our set *F* of employed feature spaces then consisted of the following combinations: *F* = {*Env*, *D*_+_*(Env)*, *Sg*, *Sg+D*_+_*(Sg)*, *Sg+Gb*, *Sg+Art1D*, *Sg+Art*}. Each feature space was downsampled to 40 Hz and z-scored prior to modeling.

### Mapping from stimulus to MEG

To perform a linear mapping from our feature spaces to the recorded MEG signals, we used ridge regression (Crosse et al., 2016) in a 6-fold nested cross-validation framework (Varoquaux et al., 2017). This allowed us to tune hyperparameters controlling the temporal extent and the amount of L2 regularisation of the ridge models as well as the amount of regularisation of the sensor covariance matrices and the coordinates of positions in source space for the beamformer spatial filters in the inner folds, yielding data-driven optimised models for each feature space, hemisphere and participant.

#### Linear model

The single-subject linear model we employed can be formulated in discrete time as:

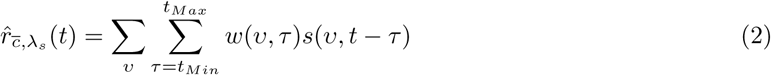

Here, 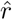 denotes the neuronal response as obtained with a spatial filter with maximum gain at the vector 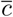 of coordinates (X, Y, Z) in source space and a regularisation of the sensor covariance matrix of λ*_s_*. Further, *s* is a representation of the stimulus in a given feature space, possibly multidimensional with dimensions *ν*. Finally, *w* describes the filter weights across these dimensions and time lags *τ* ranging from *t_Min_* to *t_Max_*, where negative values refer to samples in the future of *t* and positive values refer to samples in the past of *t*.

To obtain these filter weights, we used the following closed-form solution:

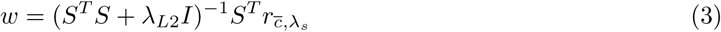

Here, S denotes the lagged time series of the stimulus representation, each column consisting to a particular combination of lags *τ* and feature dimensions *ν*, organised such that neighbouring feature dimensions populate neighbouring columns within groups of columns corresponding to time lags. The identity matrix *I* is multiplied with λ_*L*2_, a hyperparameter adjusting the amount of L2 regularisation. Larger values of λ_*L*2_ force the resulting weights *w* closer to zero and thus reduce overfitting.

For the joint feature spaces consisting of multiple subspaces, the temporal extent and L2 regularisation was optimised individually for each subspace to obtain the best possible prediction performance. This meant that the matrix *S* was constructed as the columnwise concatenation of multiple submatrices with different numbers not only of feature dimensions *ν* but also of lags *τ*. Additionally, this meant that λ_*L*2_ here was a vector instead of a scalar, with as many elements as feature spaces in the joint space. Corresponding to the concatenation of *S*, different sections of the diagonal of the identity matrix were multiplied with the dedicated regularisation parameters of the corresponding subspace.

We used an additional regularisation for the *Gb* feature space. We had observed that feature dimensions belonging to the group of fastest temporal modulation frequencies *ω* had noisy and small filter weights at long absolute temporal lags. Based on this, we concluded that the temporal extents *τ* chosen for this feature space were essentially a compromise of long optimal *τ* for feature dimensions of slow *ω* and short optimal *τ* for feature dimensions of fast *ω*. To remedy this problem, we assigned the usual *τ* to the group of slowest *ω* and added additional *τ* hyperparameters for the group of fastest *ω*. The *τ* of the central *ω* were then spaced proportionally to the mean auto-correlation times (ACT) of the corresponding groups of feature dimensions of this stimulus representation. We defined the ACT as the shortest lag where the normalised and absolute auto-correlation dropped below a value of .05. This allowed the optimisation algorithm to pick long *τ* for feature dimensions of slow *ω* and short *τ* for feature dimensions of fast *ω*.

#### Nested cross-validation and hyperparameter tuning

To make data-driven optimal choices for the range of lags *τ* defined by *t_Min_* and *t_Max_*, the amount of L2 regularisation λ*_L2_*, the coordinates in source space as well as the amount of regularisation of the sensor covariance matrices λ_*source*_, we used nested cross-validation. Specifically, this means that we split our stimulus and response data in six portions of equal durations. Two loops then subdivided the data into training, tuning and testing sets. In each iteration, an outer loop assigned each of the six portions to be the testing set. Additionally, in each iteration of the outer loop, a full run of an inner loop was performed, assigning four portions to be the training set and the remaining portion to be the tuning set. This resulted in a total of 30 different assignments of portions to different sets. With this framework, we first picked a certain combination of hyperparameters and computed the corresponding weights *w*, the elementary parameters, using the training set. The resulting filters were convolved with the stimulus of the tuning set to obtain predictions 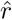 which we correlated with the observed responses *r* to obtain the tuning performance. This was repeated 200 times with different combinations of the hyperparameters.

These combinations were chosen by a recent black-box optimisation algorithm, Bayesian Adaptive Direct Search (BADS, Acerbi & Ma, 2017). BADS uses Gaussian Processes to construct a computationally cheap internal model of the multidimensional performance landscape using already available evidence and smoothness assumptions. As the computationally relatively costly linear models are evaluated across iterations, more evidence about the true performance landscape builds up which is used to update the internal model, i.e. assumptions about the smoothness and shape of the performance landscape at hyper-parameter combinations not yet evaluated. The internal model is used to update an acquisition function, whose maximum determines which combination of hyperparameters would be most informative to evaluate next in order to find the global optimum of the performance landscape. While this algorithm is not guaranteed to find the optimal combination, i.e. it is possible that it gets stuck in local optima, it has been shown to outperform other black-box optimisation algorithms on datasets typical for cognitive neuroscience (Acerbi & Ma, 2017). The values at which the hyperparameters were initiated as well as the ranges to which they were constrained are shown in table 1.

Once all iterations of an inner loop were finished, we averaged the hyperparameter choices of all inner folds. We then retrained the elementary model parameters with stimulus and response data corresponding to these averaged hyperparameters on all five possible assignments of data portions to training sets in the current outer fold. We subsequently averaged the elementary parameters across inner folds and used the resulting weights to perform a prediction on the test set of the current outer fold. This was repeated for all outer folds to obtain a number of test set predictions corresponding to the number of outer folds.

### Model comparisons

#### Bayesian Hierarchical Modeling of performances

In an initial evaluation of the encoding models, we wanted to statistically compare the predictive performance from multiple predictors, obtained from multiple participants. Similar situations often arise in neuroimaging and are usually complicated by small raw effect sizes across conditions in the presence of much larger between subject variability. A promising way to address this is provided by hierarchical models, which allow to maintain sensitivity to effects of interest in these cases. To evaluate the model performances *r* in both hemispheres *h* for each outer fold *b* of all participants *i* and focus on the differences between the feature spaces *f*, we used a Bayesian hierarchical model with a zero intercept, participant-independent and participant-specific effects for each feature space as well as effects specific to each combination of participants and folds as well as each combination of participants and hemispheres. This allowed us to assess posterior distributions of the beta estimates of the means of each level of the categorical variable feature space. To implement this model, we used the brms package (Bürkner, 2017) within the R computing environment (R Core Team, 2013). Specifically, the chosen package implements a user-friendly interface to set up Bayesian hierarchical models using stan (Stan Development Team, 2018). We used Markov chain Monte-Carlo sampling with four chains of 4000 iterations each, 1000 of which were used for their warmup. The priors for all parameters were not changed from the default values, i.e. (half)-student-*t* distributions with 3 degrees of freedom for the intercepts and standard deviations. The model can be described with the following formula:

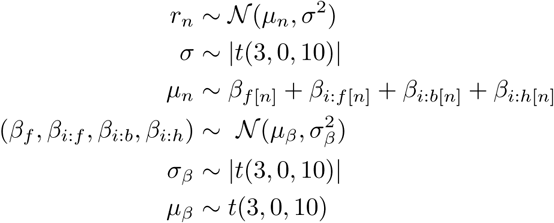

To compare the resulting posterior distributions for several parameter combinations of interest, we evaluated the corresponding directed hypotheses using the brms package: β*_f_a__* − β*_f_b__* > 0, for all possible pairwise combinations of feature spaces, and obtained the ratio of samples of the posterior distributions of differences that were in line with the hypothesis.

#### Partial Information Decomposition

Besides directly comparing the raw predictive power of models across feature spaces, we were also interested in characterising the detailed structure of predictive information carried in the different feature spaces. Since we were particularly interested in discovering to what degree the contributions of the annotated feature spaces can be explained with contributions of acoustical feature spaces, we thus asked to what degree their predictions contained the same information about the observed MEG (redundancy) or to what degree their contributions were distinct (unique information). In information theory, this is possible within the framework of Partial Information Decomposition (PID, Williams & Beer, 2012; Wibral et al., 2015) of systems of two source- and a target variable. This can be seen as a further development of the concept of interaction information (McGill, 1954) or co-information, which however conflates redundant with synergistic information and cannot resolve unique contributions.

We used a recent implementation based on common change in surprisal (*I_ccs_* Ince, 2017) which has been applied within a neuroimaging context before (Park et al., 2018). Here, the basic idea is to resolve mutual information on the local level, where it can be understood as representing the difference in surprisal a value of one variable has when the value of another is observed. Contrary to global mutual information, this change in surprisal is then a signed quantity, reflecting positive information and negative misinformation. For *I_ccs_*, only terms are considered where two conditions are met: (I) both sources have local information about the target of the same sign and (II) the local co-information of these three variables is of the same sign as the two local mutual information values. This allows to quantify contributions of the sources about the target which the sources genuinely share, i.e. redundancy. A crucial advantage of this redundancy measure as opposed to other PID implementations is that it measures the overlap at the pointwise level and therefore can be interpreted as a within sample measure of redundant prediction. Once this is done, the other quantities in PID, namely unique information and synergy, can then be inferred via a lattice structure (Williams & Beer, 2012).

We here performed PIDs for each combination of outer fold predictions of the annotated feature space with those of the acoustic feature spaces as sources and the observed responses as targets. Critically, we retrained all models with fixed hyperparameters of regularisation of sensor covariance matrices and coordinates in source space to those previously chosen as optimal in the inner folds when training the model based on the annotated feature space. This way, we gave the annotated feature space the best chances to achieve maximal unique information. To compute the respective information theoretic quantities with these continuous variables, we copula transformed the variables (Ince et al., 2017) prior to running PIDs for gaussian variables via Monte Carlo integration (Ince, 2017).

#### Phoneme-evoked dynamics

A recent study reported that epoching EEG recordings from a story-listening paradigm according to the onsets of phonemes allowed the decoding of four classes of phonemes, so-called manners of articulation, from the resulting event-related potentials (Khalighinejad et al., 2017). We aimed to firstly replicate this finding with our MEG data and secondly assess to which degree our linear encoding models could account for this phenomenon.

We computed “Phoneme-Related Fields” (PRFs) using the 34562 phoneme presentations we had previously identified in our stimulus material. For this, we mapped the set of phonemes to manners of articulation as specified by Khalighinejad et al. (2017): Plosives, fricatives, nasals and vowels. We then epoched the continuous MEG data for a time range from −.1s – +.6s around phoneme onset, binned it across epochs for each time point using four equipopulated bins and computed mutual information between the MEG data and the four manners of articulation. To ensure that we would capture the maximum effect of the MI, we delegated the choice of source positions for the left and right hemispheres as well as sensor covariance regularisations to the BADS algorithm similarly as before. However, this time we optimised the source model parameters with respect to the sum of MI of observed MEG data about the phoneme classes across time points. We then retrained our encoding models with the source model parameters fixed to these choices. Subsequently, we performed the same PRF analysis on the outer fold predictions of each feature space. We were then interested in the redundant and unique contributions of observed and predicted MEG to the MI about manners of articulation. We thus performed PIDs with observed and predicted PRFs as sources and the manners of articulation as the target, separately for each feature space, yielding phoneme-related redundancy as well as unique profiles.

## Supplemental Material

### Hyperparameter choices

To reduce the amount of a-priori choices and improve the performance of our models, we attempted to find optimal combinations of hyperparameters of our models. Since the models were non-differentiable with respect to these hyperparameters, we used a recent black-box optimisation algorithm (Acerbi & Ma, 2017) within a nested cross-validation framework. This allowed us to efficiently control the temporal extent and L2 regularisation as well as source positions and sensor covariance matrix regularisation of our participant-, hemisphere- and feature space specific models.

We were interested in the absolute amount of variance between the chosen source positions. A low degree would reflect that the black-box optimisation would converge on the same location, a high degree would suggest that either the algorithm would get stuck in local optima or a failure in the co-registration of MEG data and anatomical scans, for example caused by excessive movement by the participant. We found the overall amount of variation between the chosen source locations to be rather small (figure S1, top left). In the worst case, the source locations were scattered within a range of 3.06 cm, the median of this range was .61 cm, only slightly above the amount we allowed the participants to move in the scanner. In the best case, the range was only .23 cm. Overall, this suggests that while there was no direct mapping of feature spaces to source positions, the optimisation of the source positions tended to converge on relatively small regions within a participant.

We were also interested if our optimisation would consistently pick distinct locations in source space for different feature spaces. To examine this, we computed the silhouette index, measuring how similar the source positions across outer folds found for one feature space were compared to those found for other feature spaces. Across feature spaces and hemispheres, we found results that were mostly inconsistent across participants (figure S1, middle). Specifically, we observed participants for whom the assignment of chosen source positions to feature spaces was appropriate as reflected by silhouette indices close to 1, but also participants for whom this assignment was inappropriate as reflected by silhouette indices close to −1. In both hemispheres though, we observed positive grand averages for the *D*_+_*(Env)* feature space, reflecting that for this feature space, relatively distinct source positions tended to be found in many participants. In sum however, the results suggests that on a group level and across feature spaces, there was no clear relationship between the choices of positions in source space and the feature space used to model the MEG responses.

The choices of optimal hyperparameters for the beamformer spatial filter did not differ substantially across feature spaces (figure S1, bottom). While we observed a relatively high degree of variance in optimal choices across participants, we found the choices to be relatively consistent within one hemisphere of a participant and across feature spaces as indicated by relatively high intra-class correlation coefficients across participants with outer folds and feature spaces as different measurements for the left (.96) and right (.88) hemispheres. However, we observed a pronounced difference between left and right ACs, with a higher level of regularisation for the left AC. Here, in some cases the optimal values even bordered on the boundaries we chose for the hyperparameter, suggesting that in some cases, even higher values could have been optimal.

For the temporal extent, it can be seen that this optimisation resulted in characteristic temporal extents for each feature (sub-)space (figure S2, top). For example, for the combination of articulatory features and log-mel spectrograms the optimisation algorithm consistently found shorter temporal extents for the articulatory features than for the log-mel spectrogram. Pooled across participants, we observed very similar patterns in left and right ACs.

**Figure S1:**
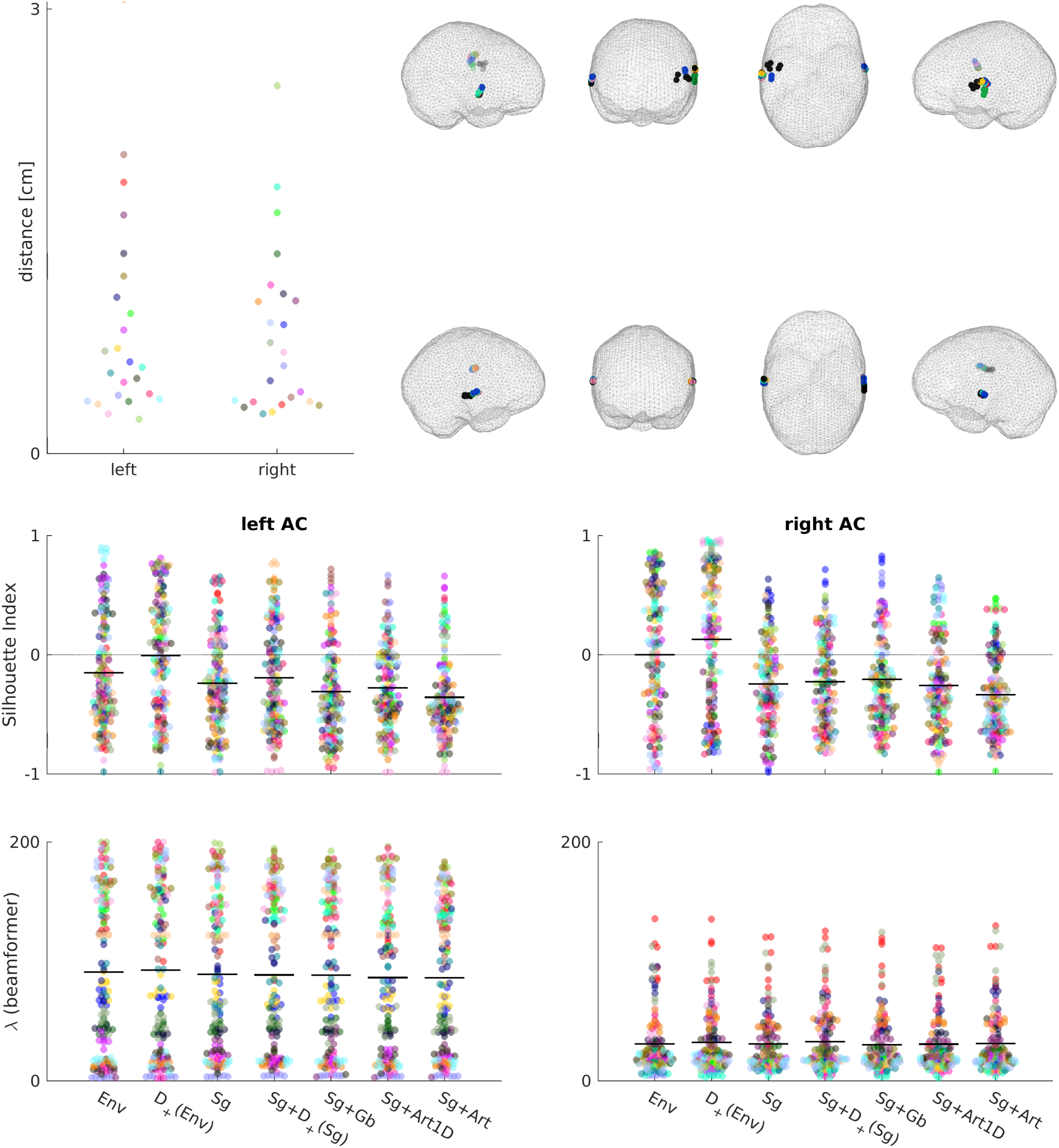
Hyperparameter choices for source model optimisation. *Top*: Chosen source positions. *Left* scatter plot shows the maximum distance between source positions used for test set prediction across all outer folds and feature spaces. Each dot is one participant in the respective hemisphere, each participant has a distinct colour. Source plots on the *right* show positions of all outer folds and feature spaces in meshes of individual brain volumes of two exemplary participants, the participant with the largest maximum distance between chosen source locations in the *top* row and the participant closest to median of maximum distance between chosen source locations in the *bottom* row. Each dot is one outer fold, colour denotes the feature space. *Middle*: Evaluation of spatial clustering of choices of source positions for different feature spaces. Each dot is one outer fold of one participant, each participant has a distinct colour. Black lines denote mean across participants and folds. *Bottom*: Choices of beamformer regularisation hyperparameters for each feature space. Each dot is one outer fold, each participant has a distinct colour. Black lines denote mean across participants and folds.

**Figure S2:**
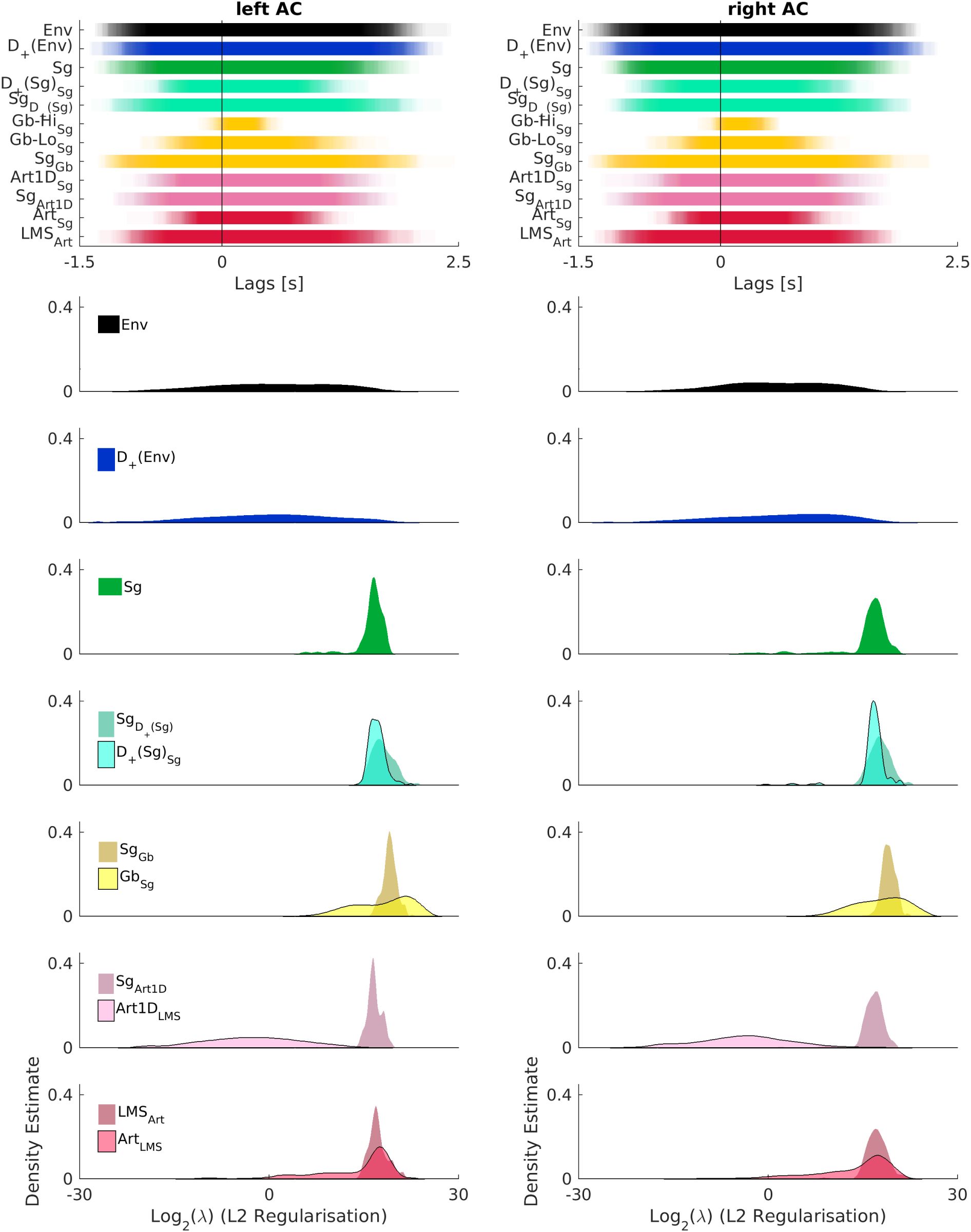
*Top*: Choices of temporal extent hyperparameters for each feature (sub-)space. Shown are averages across inner folds used for outer fold predictions, pooled across participants. Each outer fold is plotted transparently, such that the opacity codes for the number of participants and outer folds for which the temporal extent was chosen correspondingly. *Bottom*: Choices of L2 regularisation hyperparameters for each feature (sub-)space. Shown are distributions of choices averaged across inner folds used for outer fold predictions, pooled across participants.

For the L2 regularisation, it can again be seen that the optimisation found characteristic values to be optimal for each feature (sub-)space. Specifically, for lower dimensional feature (sub-)spaces the amount of L2 regularisation seemed to be less critical, yielding flat distributions. However, for higher-dimensional (sub-)spaces, a higher value of regularisation seemed to be beneficial (figure S2, bottom). This was especially the case for the combination of articulatory features and the log-mel spectrogram, for which the distributions for the two subspaces clearly differ.

**Figure S3:**
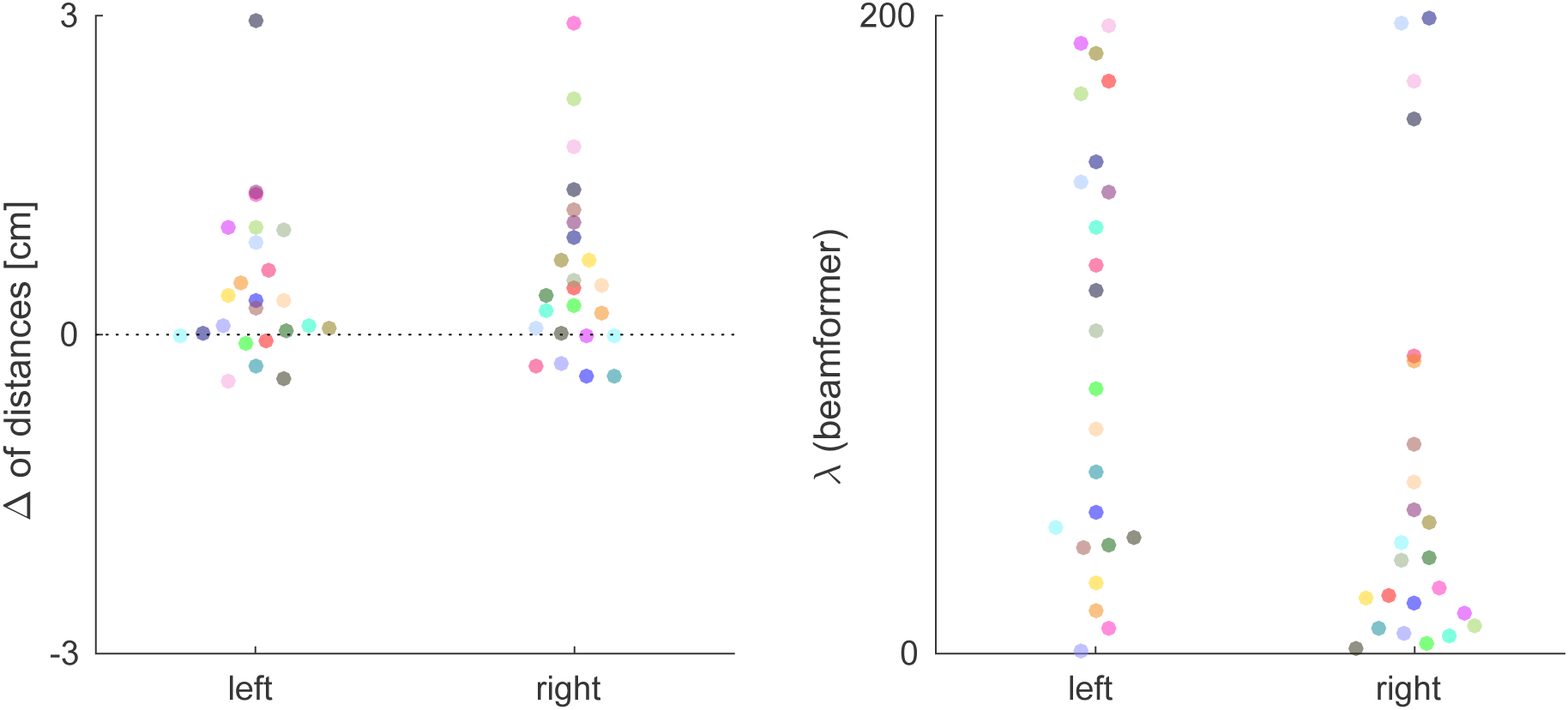
Hyperparameter choices for PRF analysis. *Left*: Maximal euclidean distances of source positions when optimised with regard to model performances across all outer folds and feature spaces subtracted from maximal euclidean distances of source positions when positions found when optimising with regard to PRF MI are included. *Right*: Results of optimising sensor covariance regularisation parameter with regard to PRF MI.

In order to provide the best possible chances for unique information of the observed MEG data when compared to the predicted MEG data in the decoding analysis, we re-optimised the source positions with respect to the point in source space with maximum information about manners of articulation, summed across time points. We recalculated the maximum distance metric used in (figure S1, top left), this time also including the positions found for optimal phoneme class decoding and plotted the difference to the previously obtained maximum differences (figure S3, left). The results reflected that still, all positions lay in STG, while for some participants, the positions found to be optimal for the PRF analysis were different from those obtained during the modeling.

Taken together, these results show that although it can not be guaranteed that the optimisation algorithm converges on global optima, it did find positions in source space confined to a relatively restricted region, sensor covariance regularisations with a high degree of within-participant similarity and distinct values of temporal extent and L2 regularisation for each feature subspace. This demonstrates that the optimisation algorithm was able to pick up on general relationships between hyperparameter choices and the performance of the encoding models. It thus also suggests that the parametrisation chosen here was sensible in that it allowed the optimisation algorithm to exploit these general relationships.

## Acknowledgments

CD is funded by the College of Science and Engineering at the University of Glasgow; JG received support from the Wellcome Trust (UK; 098433). We thank Moritz Boos, Jan-Mathijs Schoffelen, Dale Barr and Christoph Scheepers for helpful discussions.

## Author Contributions

CD, RAAI, and JG conceived of and designed the experiment. CD collected and analysed the data. CD, RAAI, and JG wrote the paper. JG acquired the financial support for the project leading to this manuscript.

